# Sustainable Memristors from Shiitake Mycelium for High-Frequency Bioelectronics

**DOI:** 10.1101/2025.07.11.664296

**Authors:** John LaRocco, Qudsia Tahmina, Ruben Petreaca, John Simonis, Justin Hill

## Abstract

Neuromorphic computing, inspired by the structure of the brain, offers advantages in parallel processing, memory storage, and energy efficiency. However, current semiconductor-based neuromorphic chips require rare-earth materials and costly fabrication processes, whereas neural organoids need complex bioreactor maintenance. This study explores shiitake (*Lentinula edodes*) fungi as a robust, sustainable alternative, exploiting its adaptive electrical signaling, which is akin to neuronal spiking. We demonstrate fungal computing via mycelial networks interfaced with electrodes, showing that fungal memristors can be grown, trained, and preserved through dehydration, retaining functionality at frequencies up to 6 kHz. Notably, shiitake has exhibited radiation resistance, suggesting its viability for aerospace applications. Our findings show that fungal computers can provide scalable, eco-friendly platforms for neuromorphic tasks, bridging bioelectronics and unconventional computing.

## Background

### Overview

Artificial intelligence and neural networks are predicted to be the largest drivers of energy demand in the near future. Embodied intelligence, which uses hardware-based local neural networks, is a possible solution. These systems require ready access to large numbers of memristors, a type of electronic device that can retain information when the power is turned off, which have shown promise in enhancing the performance of computing systems. However, the manufacturing and disposal processes of conventional memristors contribute to electronic waste and environmental degradation. This context has led to interest in creating memristors using natural, biodegradable materials such as fungi.

### Memristors

Memristor devices offer substantial advantages in robotic, industrial, and transport applications due to their unique electrical properties and ability to mimic neural functions. They can enhance various control systems, facilitate efficient information processing, and ultimately improve the overall performance of autonomous systems.

One of the key strengths of memristors is their capacity for efficient and self-adaptive in situ learning, which is critical for applications in robotics and autonomous vehicles. In memristor-based neural networks, the devices can adjust their resistance according to previous inputs, allowing for a form of analog learning that closely resembles synaptic behavior in biological systems^1^. This capability enables robots and autonomous vehicles to learn from their environment and adapt in real time, enhancing their ability to navigate complex situations effectively. Studies have found that such systems can achieve low-latency responses, which are essential for high-speed decision-making in dynamic environments^2^.

Memristors also provide the advantage of integrating memory and processing capabilities into a single device, enabling a simplified architecture for autonomous control systems^3^. For instance, in autonomous vehicles, trajectory-tracking and path-following tasks can be performed using memristor-based controllers that allow for rapid calculations and real-time adjustments to control variables^4^. This integration, especially with parallelization, helps to address the challenges posed by separate memory and processing units, which can lead to delays and increased power consumption in traditional control systems^4^.

Additionally, the resilience of memristor devices against environmental changes and their ability to operate under varying conditions make them particularly suitable for autonomous applications, such as vehicles operating in unpredictable road environments^4^. This is complemented by the precision in control that memristor-based systems can offer, which is significant for maintaining stability and performance while following desired trajectories^5^.

Moreover, the low power consumption of memristors is particularly beneficial in robotics and autonomous vehicles, where energy efficiency is paramount. Hybrid analog–digital memristor systems can minimize power usage during processing without sacrificing responsiveness, which can prolong operational time by reducing the frequency at which recharging or battery replacement is required, enhancing the feasibility of deploying such systems in mobile applications^2^.

Ultimately, the potential of memristors to emulate human-like decision-making and learning processes could be exploited to endow robotic systems and autonomous vehicles with functionalities not found in conventional control systems. The ability of memristors to perform complex computations efficiently, learn adaptively, and integrate both memory and processing into a unified approach make them a cornerstone technology for the future development of intelligent autonomous systems. However, memristors often require rare-earth minerals and expensive semiconductor foundries to produce.

### Fungal Electronics

Fungi possess innate abilities to adapt to various environmental conditions and efficiently process information through their interconnected network of hyphae. These characteristics make fungi an ideal candidate for developing sustainable computing systems. We aim to design and implement a novel fungal memristor-based computing architecture that can significantly reduce energy consumption and minimize electronic waste. We can achieve this with substantially simpler bioreactors and nutrient cultures than those required for conventional neurons and neural organoids. The unique advantages of fungal memristors stem from the biological properties of fungal materials, which distinguish them from typical inorganic or polymer alternatives^6,7^.

First, one of the main benefits of fungal memristors is their environmentally sustainable and biodegradable nature. Conventional memristors often contain transition metal oxides or silicon-based structures, whose production or disposal may pose environmental challenges^6,7^. In contrast, fungal materials are derived from organic biomass, making them both sustainable and significantly less harmful to the environment. This aligns with the increasing efforts toward developing greener electronic materials, as highlighted in prior work emphasizing the importance of sustainability in technology development^8^.

Second, fungal memristors exhibit remarkable adaptability in their electrical properties. The structural composition of fungal materials often allows for a range of conductive pathways that can form dynamically under the influence of electrical stimuli, similar to the conductive filaments formed in conventional memristors^9,10^. This adaptability can lead to enhanced performance in neuromorphic applications by facilitating variable resistance states that mimic synaptic behaviors more closely than traditional memristive materials, which often have static crystalline structures that can lead to variability problems or performance limitations at the nanoscale^11^.

Furthermore, fungal memristors may consume less power than traditional materials due to their unique electrochemical properties. Prior researchers have claimed that some organic materials, including those derived from fungi, can operate effectively at lower voltages while maintaining stable switching characteristics, a trait that is crucial for developing energy-efficient devices for portable electronics and Internet of Things (IoT) applications^12^. This can significantly extend battery life and reduce energy costs in processing and memory applications, which have become focal points in research into neuromorphic systems^13^.

Finally, the natural composition and multicellularity of fungal materials can lead to more naturalistic models for neural networks. Because these materials are subject to biological processes, they may inherently incorporate characteristics that resemble biological neuronal networks, including plasticity and memory capabilities that could evolve with usage. This biological mimicry could strengthen the development of more advanced artificial neural networks, enabling applications such as adaptive learning systems and intelligent sensor networks^14^.

### Fungus Types

The potential use of shiitake (*Lentinula edodes*) and button mushrooms (*Agaricus bisporus*) as organic memristors is an emerging area of research that exploits the unique properties of these fungi. Memristors, which are non-volatile memory devices that retain information even without power, can benefit from the porous structures and electrical properties of organic materials derived from mushrooms.

Shiitake mushrooms have been shown to possess a hierarchically porous carbon structure when activated. This porous structure can enhance the electrochemical performance of devices, making them suitable candidates for use in energy storage systems, including supercapacitors and potentially memristors^15^. The creation of highly conductive carbon materials from shiitake has been demonstrated, suggesting that these materials could be engineered to exhibit memristive behavior^16^. Shiitake-derived carbon is a sustainable alternative to traditional materials and can enhance the performance of electronic devices due to its unique structural properties.

Button mushrooms have also shown significant potential in this context. Research has indicated that their porosity can be exploited to create materials with high surface areas, which are essential for the development of efficient electronic components^17^. The synthesis of carbon composites from button mushrooms has been explored, revealing their ability to function effectively in energy storage applications^17^. Furthermore, the integration of button mushrooms into electronic systems has been investigated, demonstrating their potential as substrates for electronic devices^18^.

In addition to their structural properties, the unique biological characteristics of fungi, including their ability to interact with various chemical compounds, can be harnessed to develop novel sensing technologies. For instance, electronic noses have been developed that use mushroom extracts to detect volatile compounds; these could be adapted for use in electronic devices that require environmental sensing capabilities^19,20^. This intersection of biology and electronics opens new avenues for creating multifunctional devices that incorporate the sensory capabilities of mushrooms.

### Radiation Resistance and Resilience

The radiation resistance of shiitake mushrooms has been studied primarily in terms of their ability to withstand and possibly derive benefits from exposure to ionizing radiation. This resistance can be attributed to several biochemical and physiological attributes. A possible factor is lentinan, a polysaccharide found in the cell walls of shiitake. Lentinan provides structural integrity and exhibits immunomodulatory effects that may enhance the mushroom’s ability to respond to environmental stresses, including radiation exposure. Although some research suggests that lentinan possesses properties that may help mitigate oxidative stress^21^, studies directly linking lentinan to radiation resistance in shiitake mushrooms are limited.

Shiitake mushrooms have also shown a notable ability to adapt to their environmental conditions, including variable radiation levels. Studies involving fungi in space research indicate that certain fungi can enhance their survival through morphological changes or increased melanin production in response to radiation^22^. This research has not specifically addressed shiitake, but the general adaptability observed in fungi suggests that this species could respond similarly to such conditions.

Another example of the resilience of shiitake mushrooms is their ability to maintain their nutritional and bioactive qualities after irradiation. For example, they retain essential nutrients and bioactive compounds even after exposure to ultraviolet radiation^23^. The high content of ergosterol, a precursor to vitamin D, found in shiitake mushrooms, reinforces their potential for beneficial outcomes following exposure to radiation, as this compound can be converted into vitamin D_2_ when subjected to ultraviolet light^24^.

Lastly, shiitake mushrooms can be considered in the development of dietary supplements or functional foods that might serve a broader purpose in radioprotection. Their multirole efficacy as a food source and electrical component emphasizes a sustainable approach to utilizing biological entities that can withstand environmental stresses, including radiation. This is especially relevant in aerospace and exploration contexts, where promoting health in astronauts could reduce the risks associated with increased radiation exposure during missions^22^. Lastly, shiitake mushrooms can withstand environmental stresses, including radiation, while remaining safe for human consumption.

In summary, the radiation resistance of shiitake mushrooms is linked to the presence of protective compounds like lentinan and their ability to adapt morphologically. These factors contribute to understanding their survival strategies and suggest potential applications in areas where radiation exposure is a significant concern.

## Methods

### Hyphal Cultivation

Due to the financial and environmental constraints of this project, all memristors fabricated during experimentation were composed exclusively of low-cost, organic materials. Drawing from prior research, materials such as biocompatible composites^25^ were identified as viable candidates for device construction and programming due to their biodegradability and compatibility with fungal growth.

The initial phase of experimentation focused on the cultivation of fungal hyphae within the selected organic growth media. Nine samples were prepared in standard polycarbonate petri dishes. The growth conditions were carefully maintained to promote optimal fungal development, with a controlled temperature range of 20–22°C, a relative humidity of 70%, and mixed light exposure to replicate natural terrestrial conditions. The nutrient substrate consisted of a mixture of farro seed, wheat germ, and hay, selected for their organic composition and ability to support robust fungal growth. Each sample was inoculated with spores or mycelial plugs of shiitake.

As shown in Figure 1, the samples were observed and documented biweekly to track their growth consistency and morphological development. Observations such as hyphal density, surface coverage, and color changes were recorded in a structured laboratory logbook. The log included timestamps and qualitative notes, enabling consistent comparison across samples and time points.

**Figure 1.**
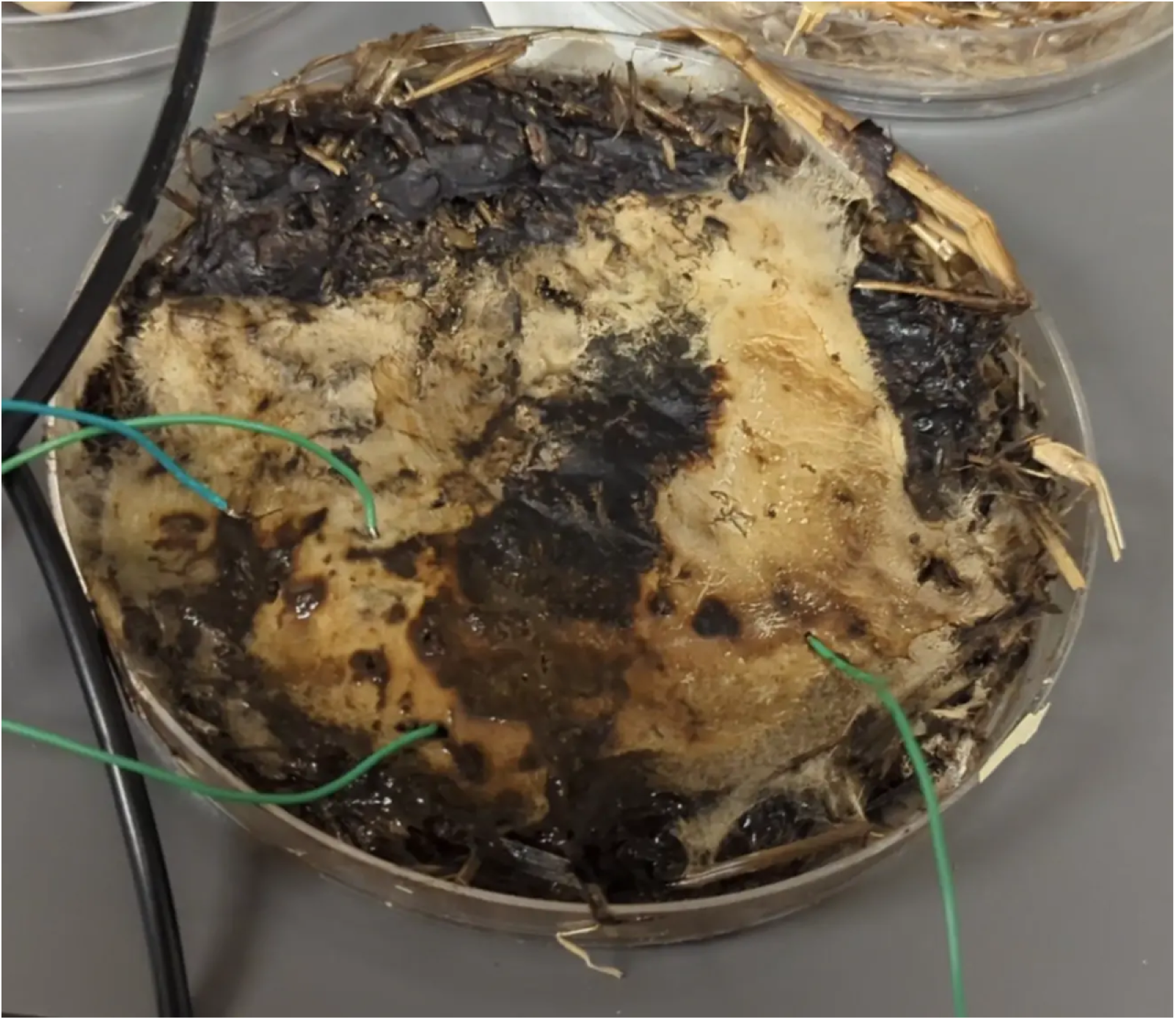
Fungal sample with probe points. Each sample grew a mycelia network that was connected to conventional electronics.

### Drying and Rehydration Process

Once full hyphal coverage and structural maturation were achieved when the petri dish was covered, the samples were transitioned to a drying phase. The petri dishes were placed in a well-ventilated area under direct sunlight for approximately seven days, ensuring uniform dehydration. The samples were rotated periodically to avoid uneven hardening. As claimed by prior work, this process transformed the fungal matrix into a rigid, disk-like structure while retaining its overall shape and connectivity.^27^

Prior to testing, the samples were rehydrated using a fine mist of aerosolized deionized water. This step restored the required level of conductivity without introducing bulk moisture that could alter their mechanical integrity.

### Electrical Characterization

Electrical testing protocols were designed in accordance with theoretical memristors. ^6,7^ An alternating current (AC) was applied to each sample, and the corresponding current–voltage (I–V) characteristics were measured using a digital oscilloscope.

To extract accurate current values, a known shunt resistor was placed in series with each sample. As shown in Figure 2, voltage readings were captured across both the sample and the resistor, with Channel 1 of the oscilloscope measuring the input voltage and Channel 2 capturing the voltage drop across the shunt resistor. Current values were then calculated using Kirchhoff’s current law, allowing derivation of the I–V characteristics from voltage differentials. All waveform data were exported in comma-separated values (CSV) format for subsequent digital analysis and visualization.

**Figure 2.**
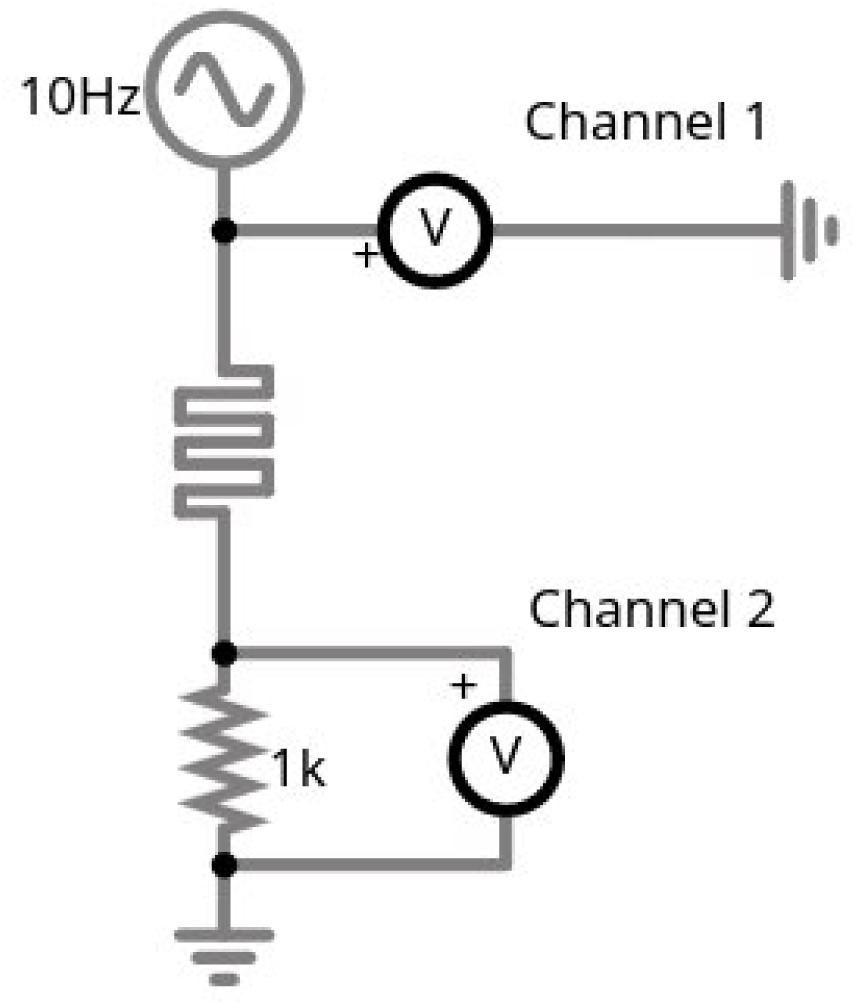
Example theoretical test circuit. The test circuit evaluated the memristive properties of each sample.

To investigate the memristive behavior of the samples thoroughly, voltage sweeps were conducted using both square and sinusoidal waveforms. Square waves were employed to detect sharp threshold-based resistance changes, whereas sinusoidal inputs provided insight into more subtle, continuous mem-fractive behaviors. This dual approach enabled the identification of hysteresis loops in the I–V curves, a key signature of memristor functionality.

### Hypothesis

The general testing setup, informed by prior literature, can indicate whether the fungal samples are capable of exhibiting memristive behavior. If present, this behavior would manifest as a characteristic pinched hysteresis loop in the I–V curves, typically intersecting at or near the origin, a well-established signature of memristive systems. This study hypothesized that such a response would emerge under specific combinations of voltage amplitude and input frequency.

### Volatile Memory Testing

After experimentally confirming that the fungal samples exhibited memristive behavior, a specialized electronic circuit was designed and implemented to investigate the volatile memory characteristics of two fungal samples in series. To parallel prior work, the configuration and layout of this testing circuit are presented in Figure 3.^6^

**Figure 3.**
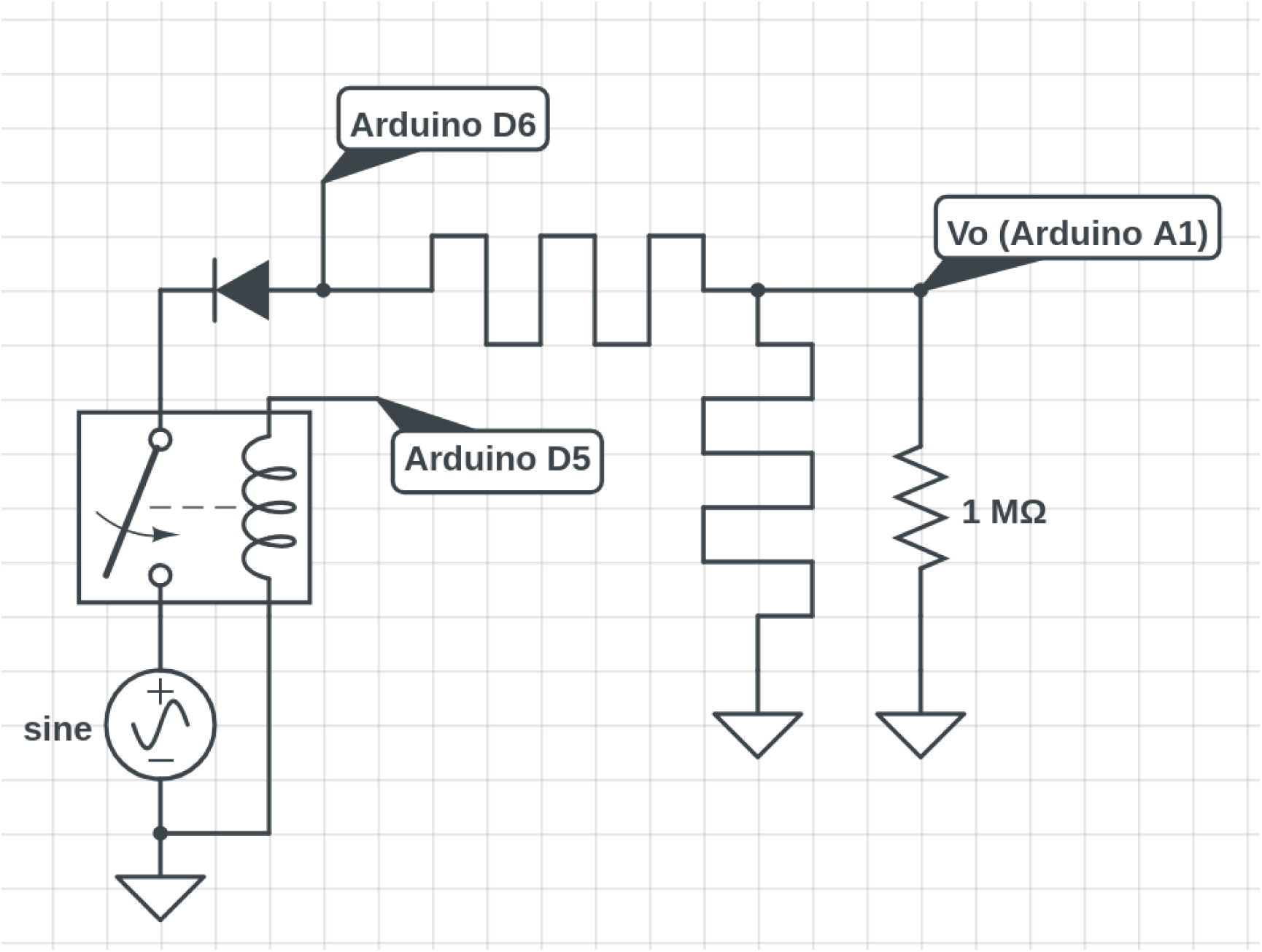
Theoretical volatile memory circuit. The samples were evaluated using this model.

Comparable to prior work in memristive computing, the volatile memory circuit employed an Arduino UNO microcontroller development board and a voltage divider consisting of two memristive elements. ^6,7^ Given the polarized nature of memristors, the circuit was designed to allow setting a voltage of opposite polarity to that used during read operations. Both voltages used were approximately 5 V. The Arduino UNO cyclically applied a high signal to a relay containing a half-rectified sine wave through one of its digital output pins when reading the memristor bridge, thereby charging the divider. This process induced an asymmetry in resistance, with the memristor closest to the input experiencing a reduction in resistance, while the output-side memristor exhibited an increase. The voltage across the divider was subsequently read using an analog input pin, and another digital pin was used to run 5 V across the divider. The Arduino interpreted the stored state as “on” only when the measured voltage exceeded a predefined threshold, effectively enabling volatile memory detection based on the transient resistance states of the memristors. Results were repeated for each sample. The physical implementation of this circuit is shown in Figure 4.

**Figure 4.**
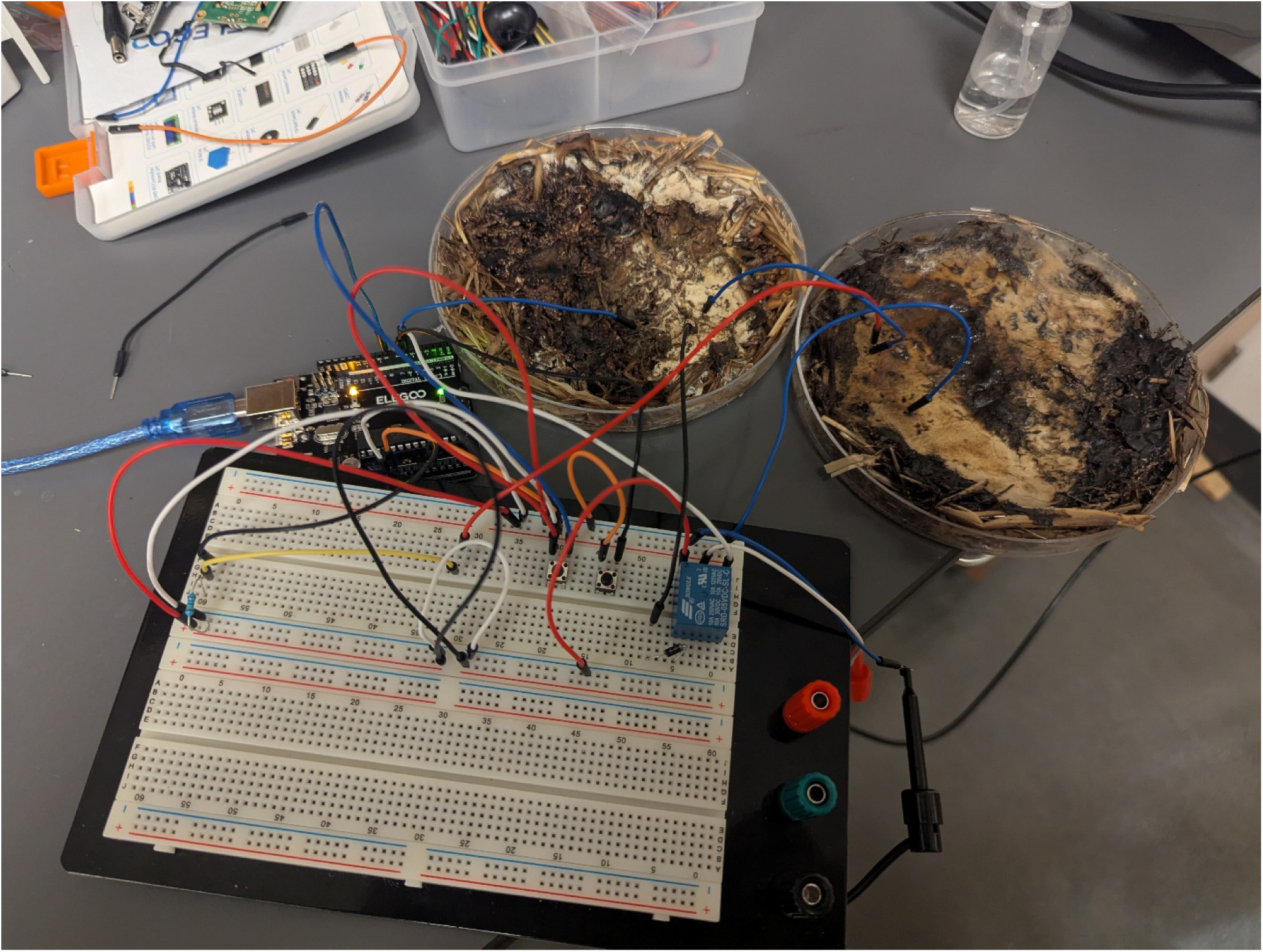
Wired samples. Volatile memory circuit implemented with fungal memristors.

## Results

### Overview

The fungal memristors were tested across a range of voltages, waveforms, and frequencies. The tested cases were as follows:

1. 200 mV_pp_ at 100 Hz, square wave
2. 200 mV_pp_ at 200 Hz, square wave
3. 20 V_pp_ at 200 Hz, square wave
4. 1 V_pp_ at 200 Hz, square wave
5. 1 V_pp_ at 200 Hz, sine wave
6. 1 V_pp_ at 100 Hz, sine wave
7. 1 V_pp_ at 25 Hz sine wave
8. 1 V_pp_ at 50 Hz, sine wave
9. 1 V_pp_ at 10 Hz, sine wave
10. 5 V_pp_ at 10 Hz, sine wave

### Voltage Testing

The first five tests were conducted to determine which voltage amplitude produced the most favorable memristive response. These initial trials revealed that a 1 V peak-to-peak (V_pp_) signal yielded the most consistent and measurable results. As outlined in the Methods section, the first four of these tests were performed using a square wave input.

### Frequency Testing

After identifying 1 V_pp_ as the optimal input voltage during the initial square wave tests (Tests 1– 4), the waveform was switched to a sine wave for further analysis (Tests 5–10). The objective of this phase was to identify the frequency at which memristive behavior, specifically a pinched hysteresis loop, became apparent.

Figures 5 through 10 show the output of Tests 2 through 7. The frequency was gradually reduced until a crossing near the origin was first observed, as shown in Figure 10. To ensure this result was not an outlier caused by overshooting the ideal frequency, the test was repeated at a slightly higher frequency (50 Hz, Test 8, shown in Figure 11). This confirmed that the optimal response occurred below 25 Hz.

Subsequently, the frequency was decreased to 10 Hz (Test 9, shown in Figure 12), which produced a clear crossing in the I–V curve near the −0.4 V region. To enhance the visibility of this behavior, the voltage was increased to 5 V_pp_, resulting in a more pronounced memristive signature (Test 10). Figure 13 illustrates this result, displaying a nearly ideal pinched hysteresis loop indicative of memristor functionality.

### Volatile Memory Experiment

The memristor voltage divider was tested by applying a 5 V_pp_ sinusoidal signal to the memristors for approximately 0.01–0.1 milliseconds. This signal was delivered via a relay triggered by digital pin 6 of the Arduino UNO. Following this brief stimulation period, the sinusoidal input was disabled, and digital pin 5 was activated to initiate the read phase. Analog voltage measurements were then acquired through the A1 analog input pin. To minimize the effects of floating voltages, a 1 MΩ pull-down resistor was connected to this pin. Voltage readings were recorded for approximately 0.1–0.10 milliseconds before the cycle repeated, allowing for rapid and continuous testing of the memristive behavior.

The results were transmitted over a serial communication interface at a baud rate of 57,600 and captured as raw text files for analysis. The data were post-processed and visualized using a custom Python script based on the matplotlib library, enabling clear identification of memory retention patterns and resistance state changes across successive cycles. The results are shown in Figures 5-18.

**Figure 5.**
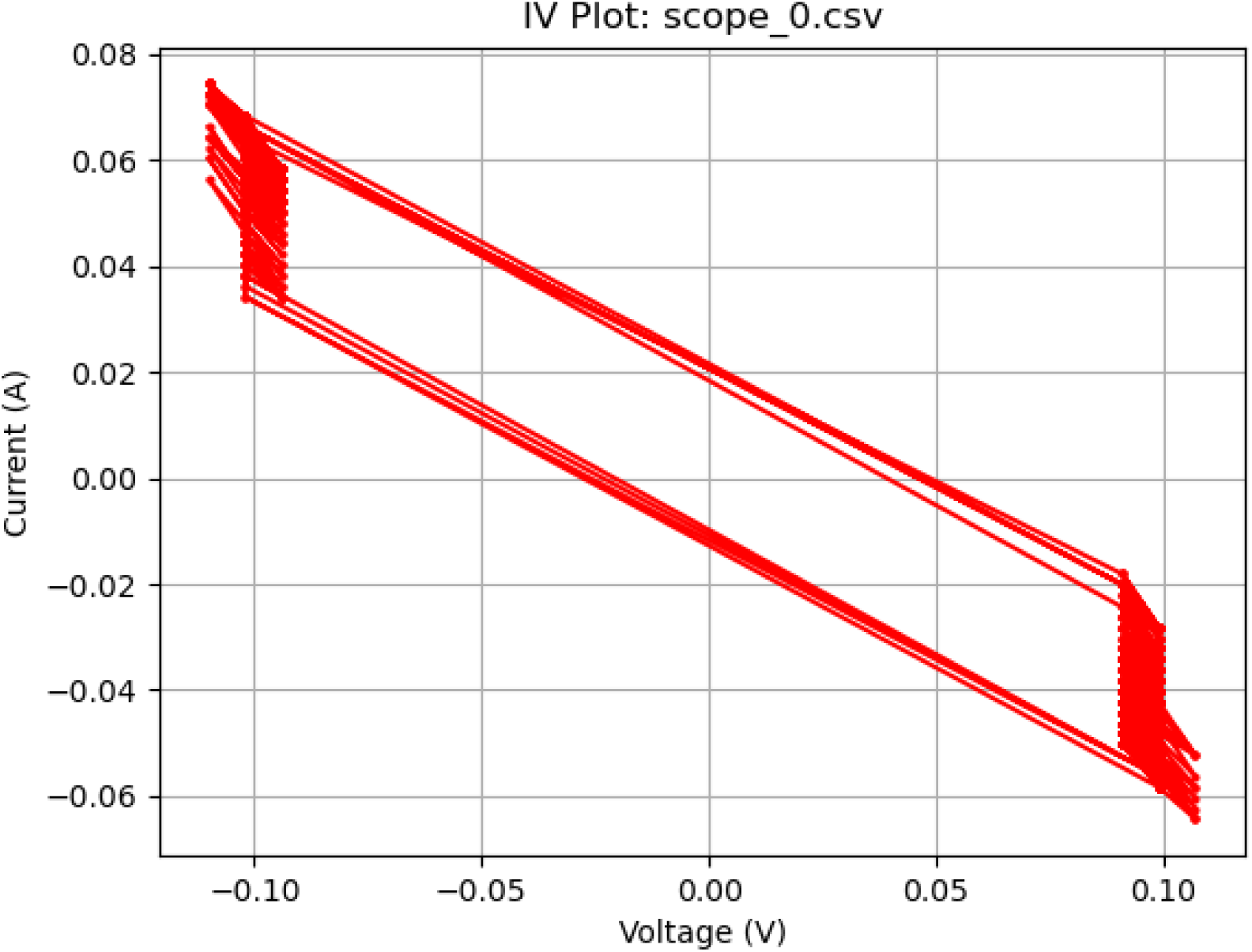
Memcapacitive activity. Plot of a 200 mV_pp_ square wave at 200 Hz displaying memcapacitive behavior.

**Figure 6.**
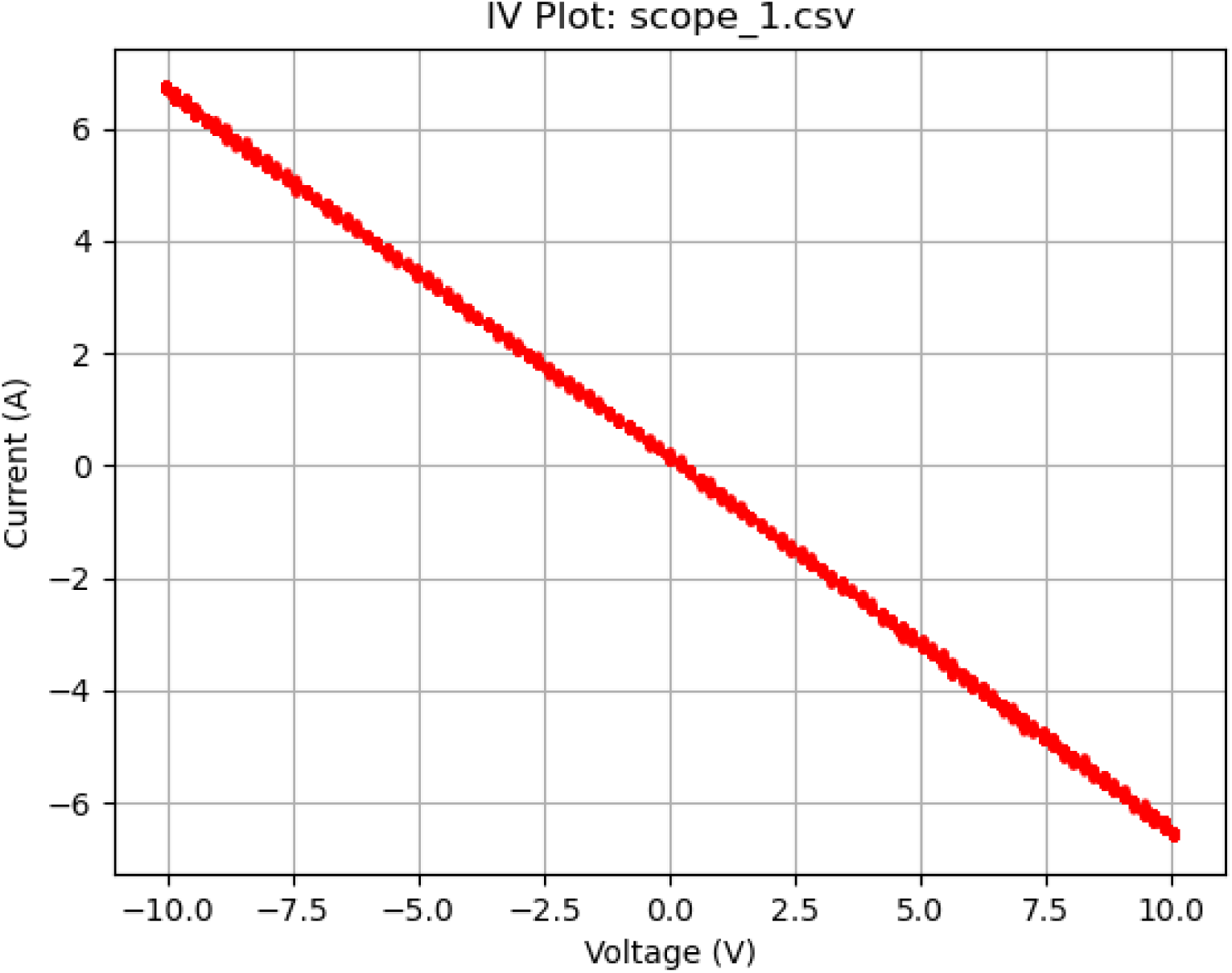
Resistive activity. Plot of a 20 V_pp_ square wave at 200 Hz displaying resistive behavior.

**Figure 7.**
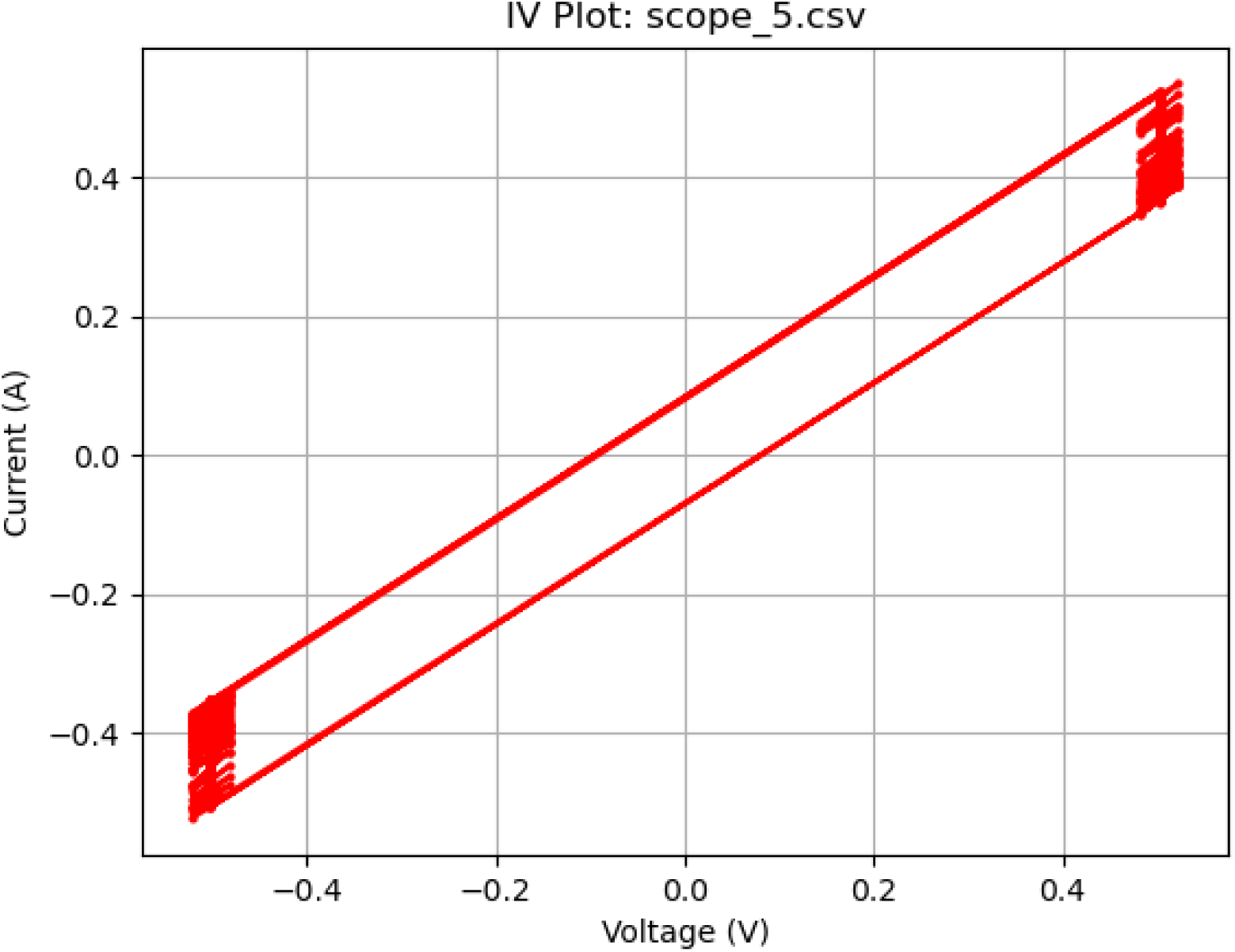
Memcapcitive activity at 200 Hz. Plot of a 1 V_pp_ square wave at 200 Hz displaying memcapacitive behavior.

**Figure 8.**
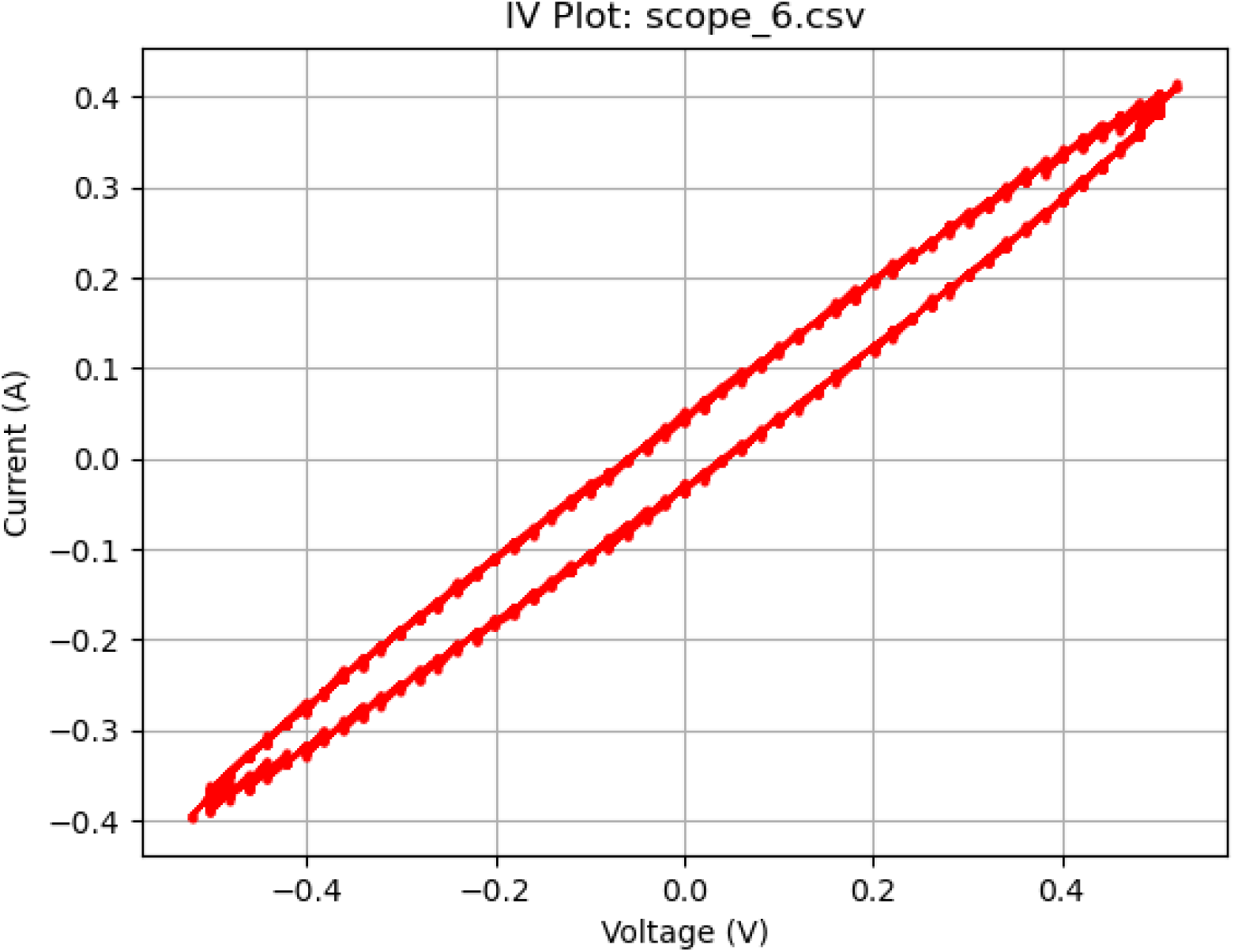
Further memcapacity activity at 200 Hz. Plot of a 1 V_pp_ sine wave at 200 Hz displaying memcapacitive behavior.

**Figure 9.**
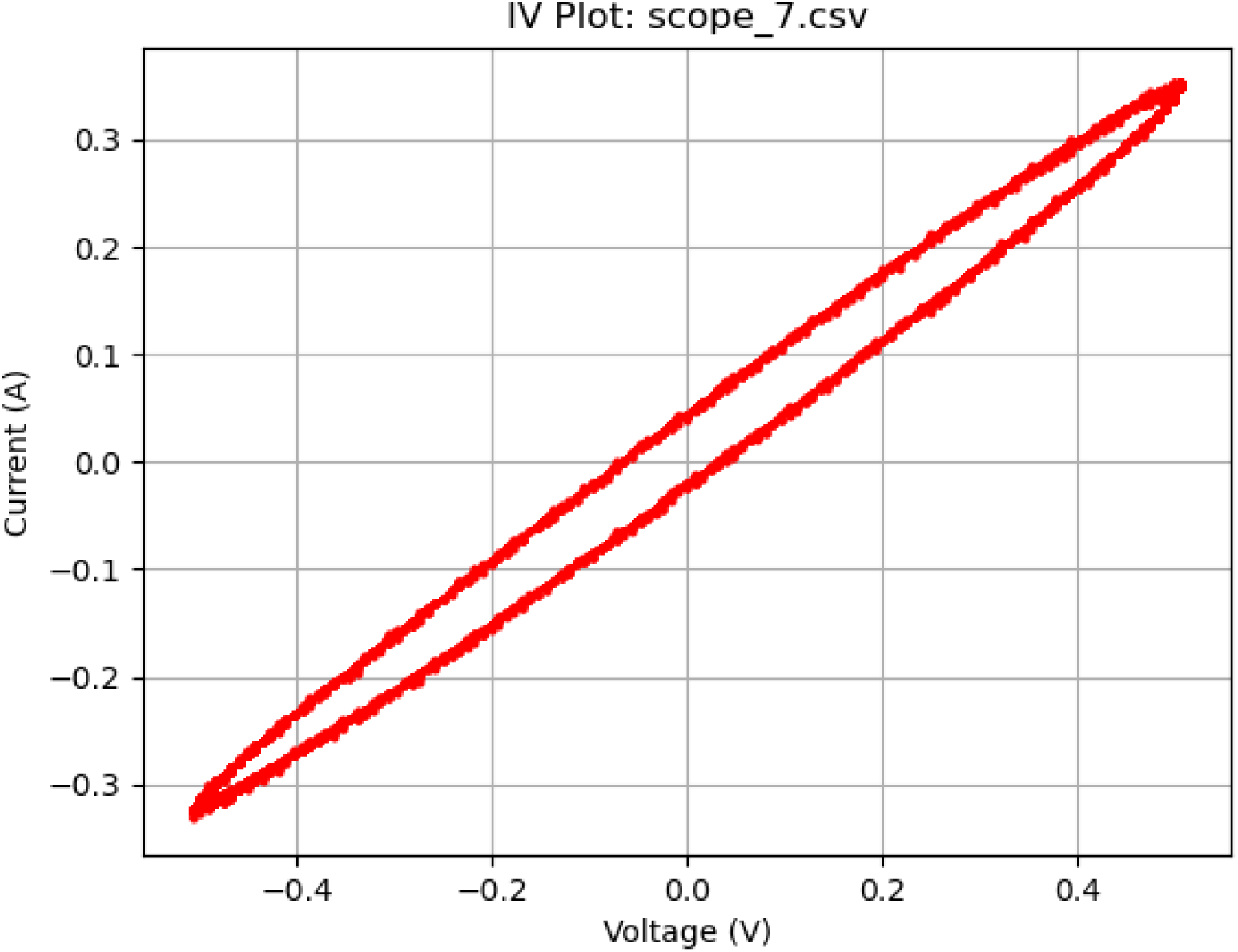
Memcapcitive activity at 100 Hz. Plot of a 1 V_pp_ sine wave at 100 Hz displaying memcapacitive behavior.

**Figure 10.**
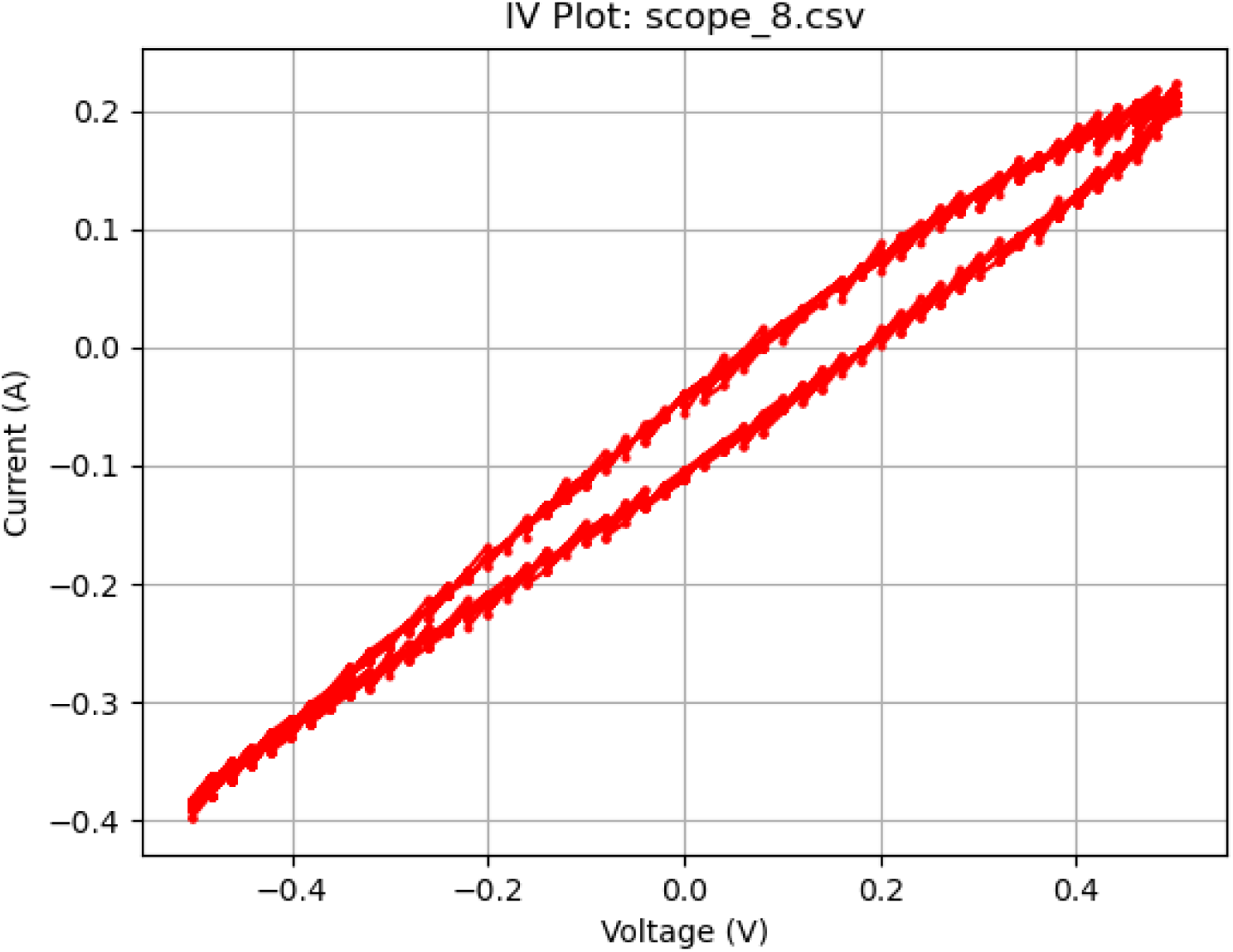
Memresistive activity at 25 Hz. Plot of a 1 V_pp_ sine wave at 25 Hz displaying memristive behavior.

**Figure 11.**
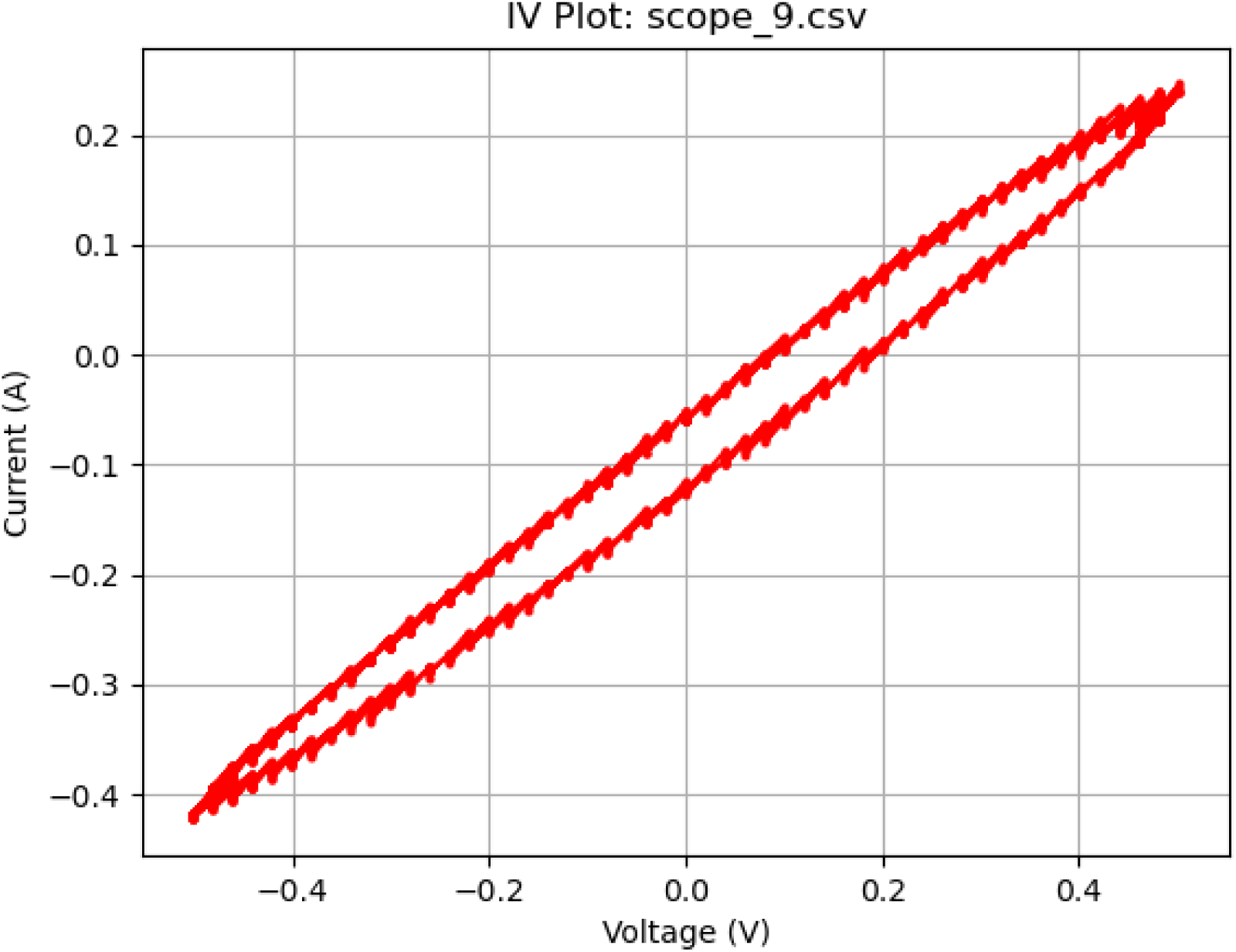
Memresistive activity at 50 Hz. Plot of a 1 V_pp_ sine wave at 50 Hz displaying memristive behavior.

**Figure 12.**
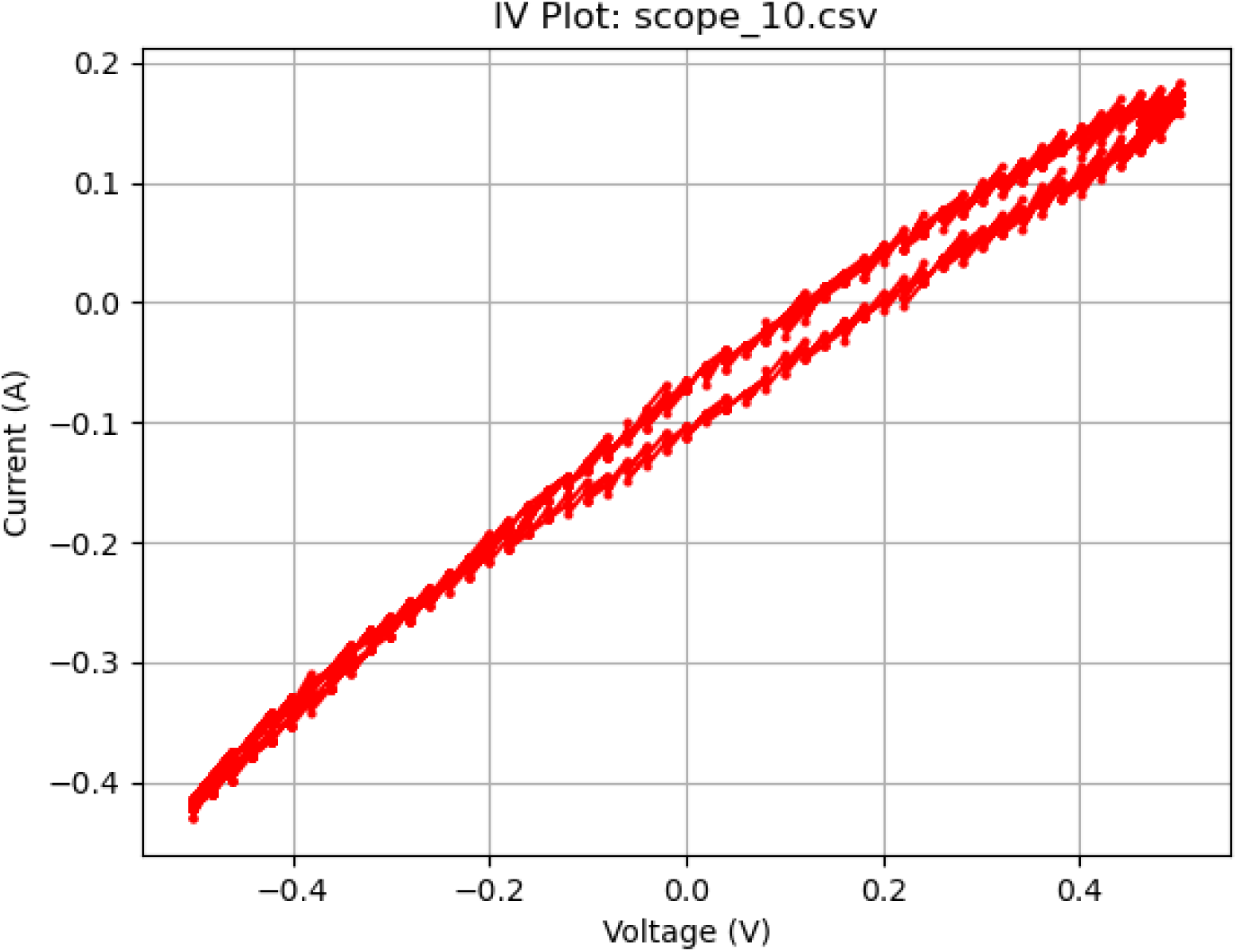
Increasingly ideal memresistive activity at 50 Hz. Plot of a 1 V_pp_ sine wave at 10 Hz displaying memristive behavior.

**Figure 13.**
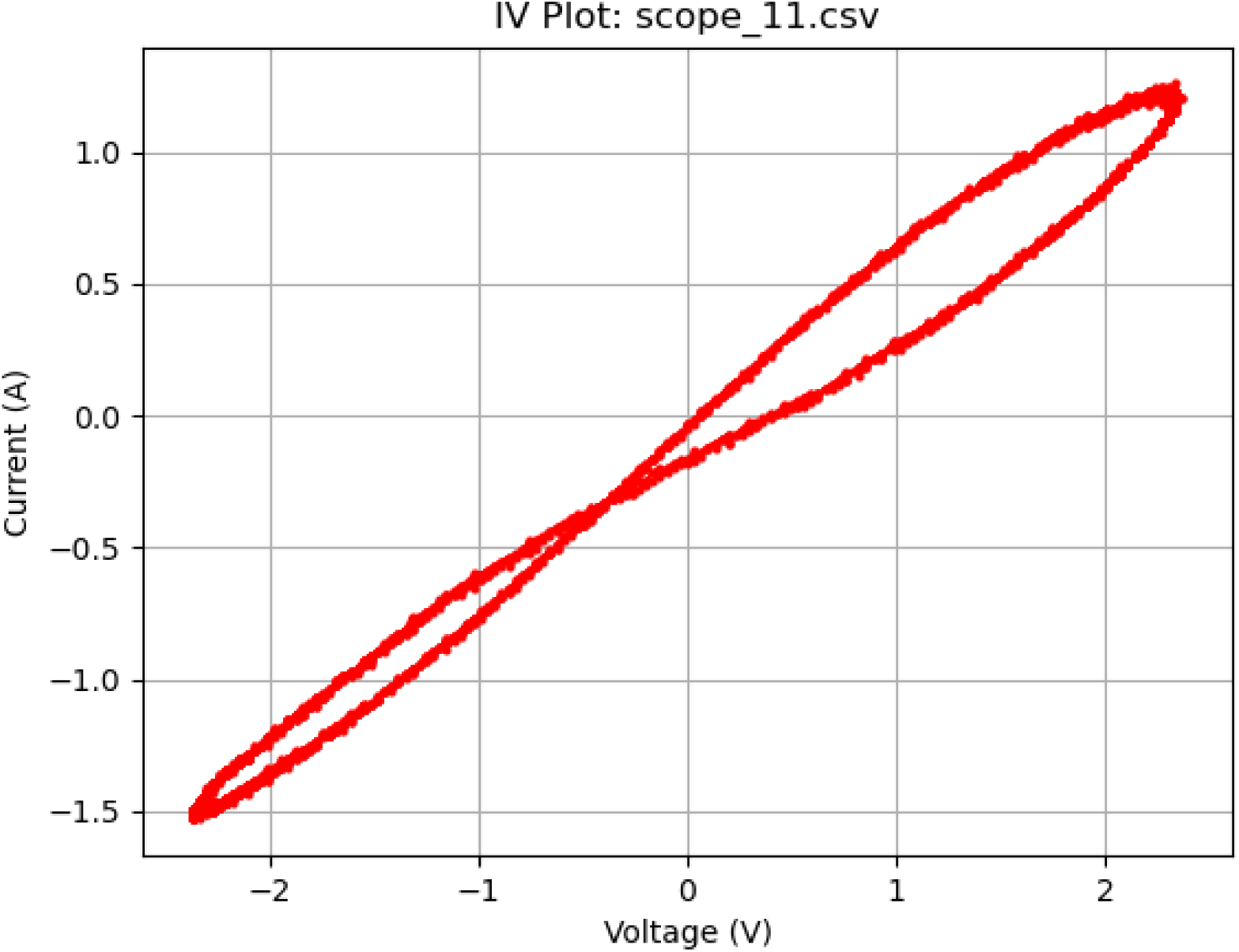
Near ideal memristive behavior. Plot of a 5 V_pp_ sine wave at 10 Hz displaying near-ideal memristive behavior.

**Figure 14.**
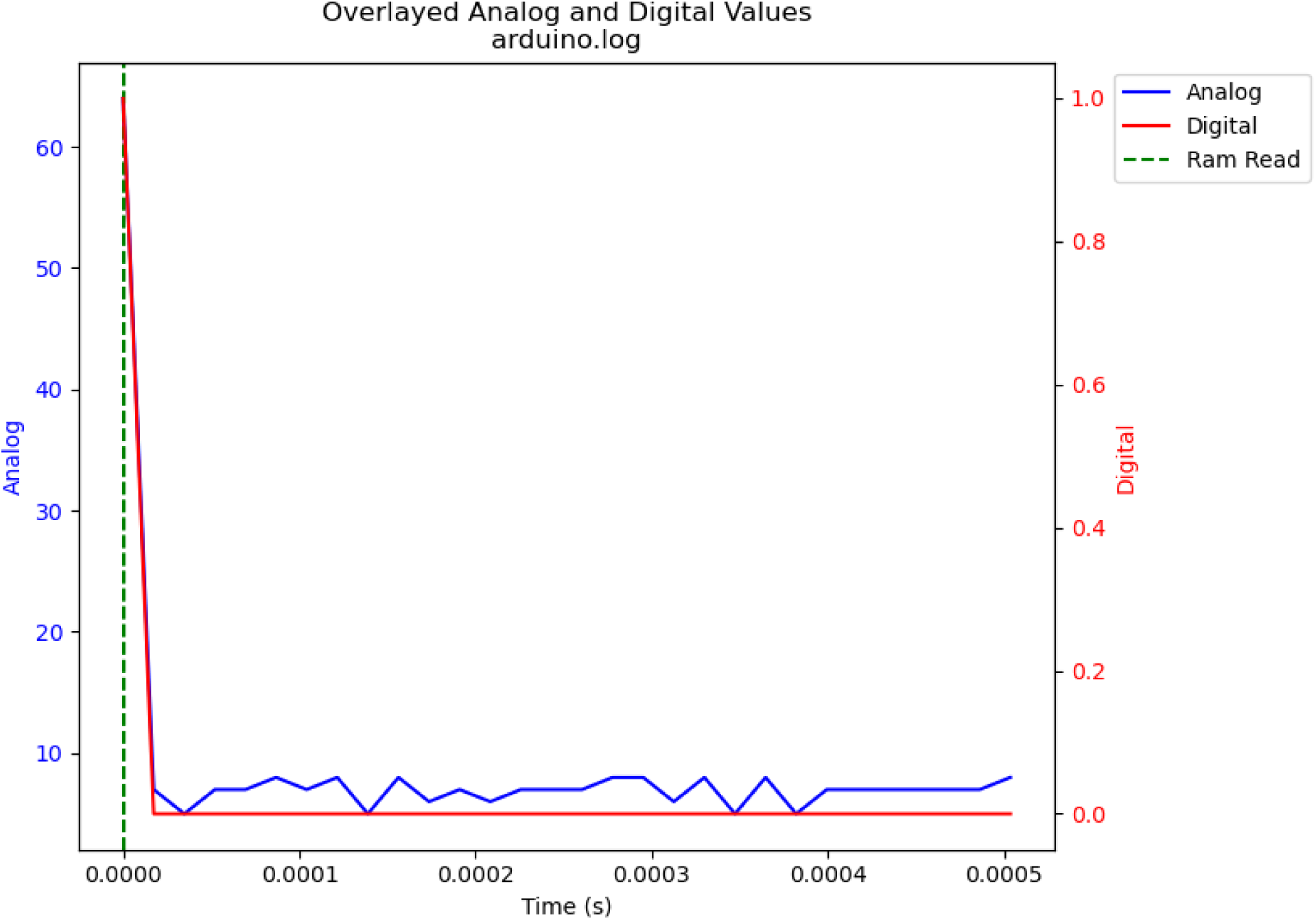
First memory test. Single write and read over volatile memory.

**Figure 15.**
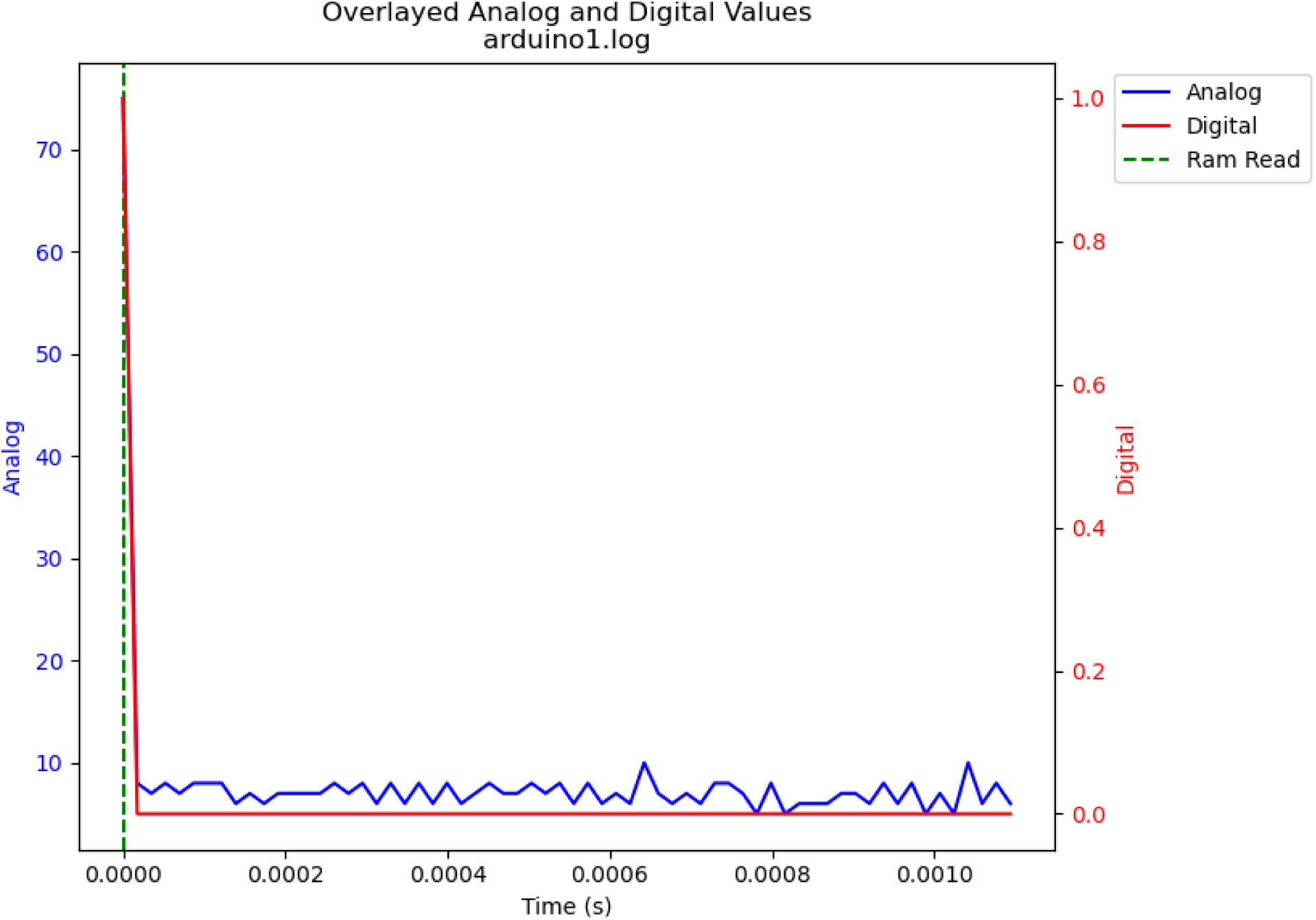
Second single memory test. Another single write and read over volatile memory.

**Figure 16.**
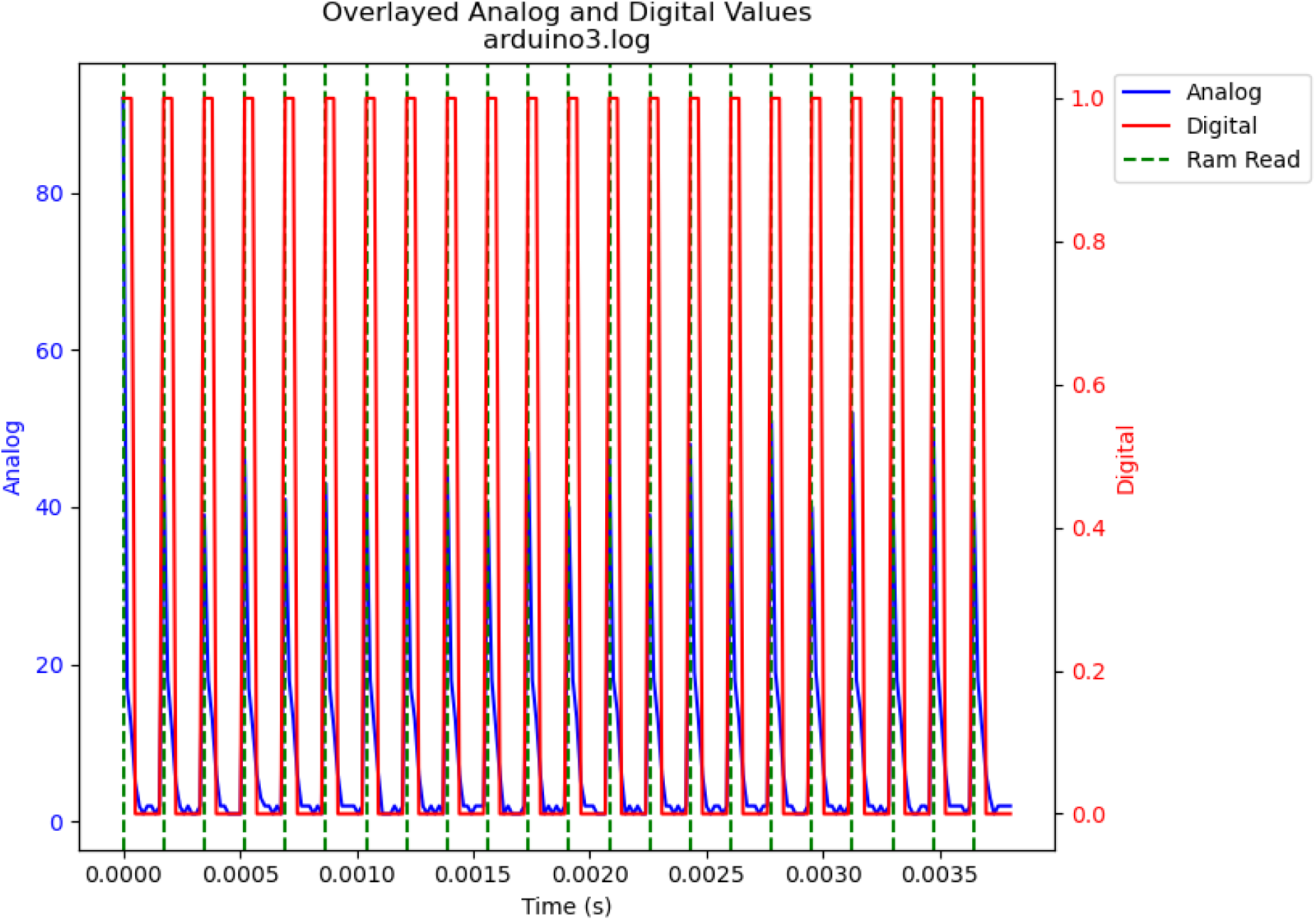
Cyclic memory test. Cyclical writing and reading over the fungal volatile memory.

**Figure 17.**
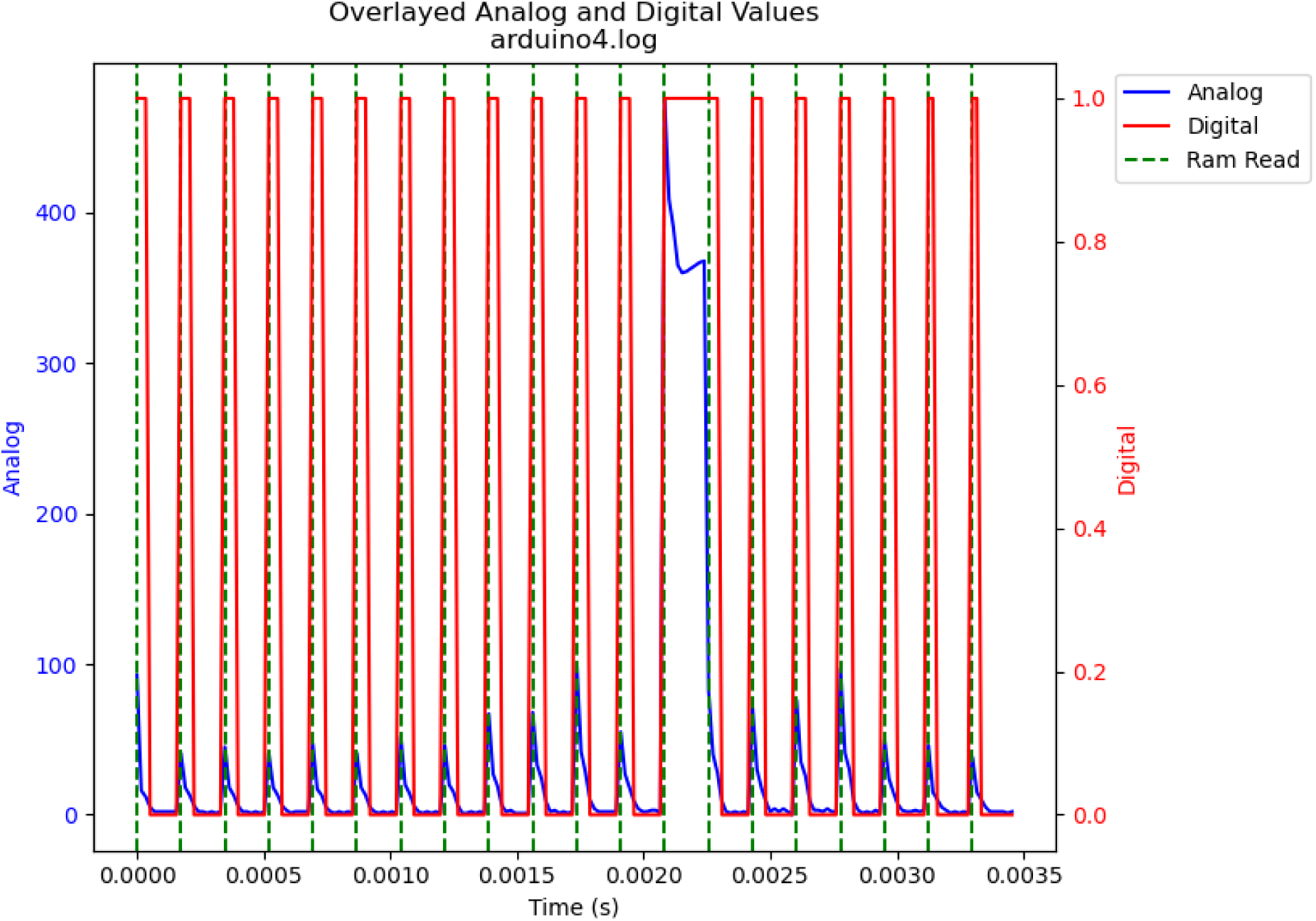
Rapid cycle memory test. Additional cyclical writing and reading over the fungal volatile memory.

**Figure 18.**
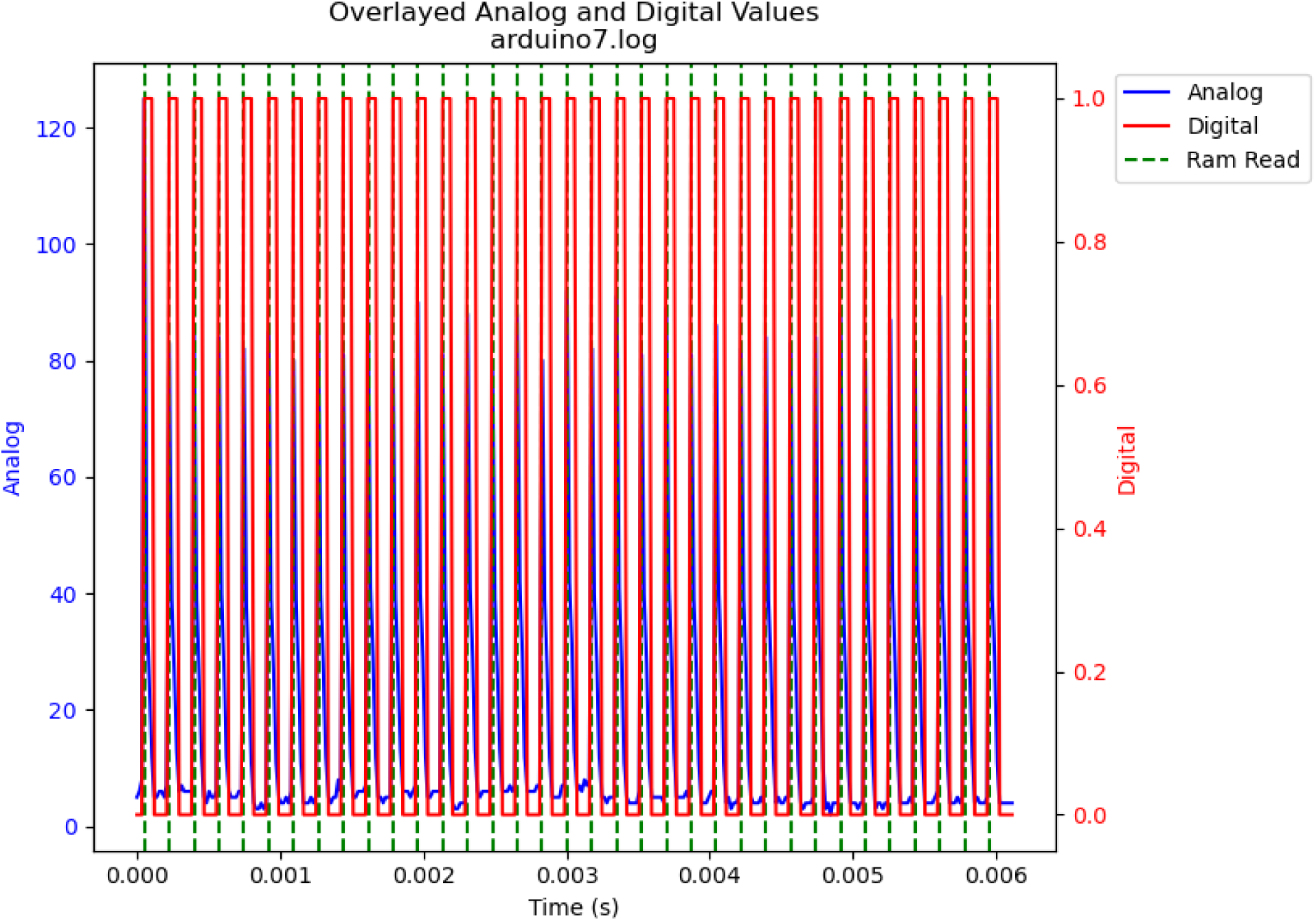
Stressed memory test. Extreme cyclical writing and reading from volatile memory.

The process and results can be summarized as follows:

- Tests 1–5: The voltage amplitude was optimized using square waves.
- Tests 5–6: The waveform was changed from square to sine.
- Tests 6–10: Frequency sweeps were performed using sine waves to identify memristive crossing.
- Test 11: The voltage range was expanded at 10 Hz (5 V_pp_) to enhance the response. This showed behavior close to that of an ideal memristor.
- Test 1: Notably, this showed consistent linear behavior, indicating resistive behavior.
- Volatile Tests 1–2: Single read and write operations were performed across the memristor voltage divider.
- Volatile Test 3: Continuous read and write operations were performed across the memristor voltage divider.

## Discussion

### Overview

Using low-cost materials, shiitake mushrooms were cultured into ideal memristors. Ideal and non-ideal memristive properties have been observed previously in fungi, but these required far more complex interfacing methods^26^. Although several techniques have been proposed to preserve fungal samples, we have obtained experimental validation that dehydration can preserve the observed properties in a previously “programmed” sample^27^. Ideal memristor properties are observed at lower frequencies, but potential latencies can be offset through massive parallelization, as in nature^26^. As with prior work on fungal memristors, the mycelial structure contains capacitive, memfractive, and memristive proteins.^25,26^ The observed rapid switching speed of 6000 Hz, low energy consumption, light weight, and radiation resistance all make fungal memristors attractive for edge computing, aerospace, and embedded firmware applications. Unlike expensive conventional memristors, culturing fungal memristors does not require large facilities or rare minerals. The process can be scaled to grow larger systems, which can be programmed and preserved for long-term use, at a low cost.

### Limitations

The current study had limitations due to its relatively short timescale of less than two months. Other researchers have documented memristive properties in mycelial materials, but their studies have also focused on short-term performance^26^. Another limitation was that only single, relatively bulky samples were prepared. To truly compete with conventional devices at the microscale and below, memristors would need to be far smaller^7,8,11^. However, the development of these devices is at an early stage, and they could eventually be miniaturized, especially with improved cultivation techniques. Complications associated with the growth media were not explored, although prior research has found that fungi are quite robust to varying conditions^26^.

### Future Work

Although fungal memristors can be produced at a low cost, certain aspects of the process could be optimized further. First, cultivation techniques could be improved using 3D-printed templates and structures that shape the shiitake mushroom into a desired geometry. Second, programming could be facilitated by adding electrical contacts into a 3D-printed cultivation structure. Finally, long-term use would necessitate preservation, which could involve a variety of techniques: dehydration, desiccation, freeze-drying, certain hydrogels, and special coatings^27^. A combination of these techniques could enable the development of fast, radiation-resistant, and low-energy memristors grown from low-cost organic materials. The future of computing could be fungal.

## Conclusions

The fabrication of existing semiconductor memristors requires rare-earth minerals and large facilities. Culturing delicate neural organoids requires a complex chemical environment to be maintained in a bioreactor. Fungal computing may provide a robust and accessible alternative. Fungal systems have lower power requirements, lighter weights, faster switching speeds, and lower industrial overheads than conventional devices. In this study, fungal memristors were fabricated, programmed, and tested using shiitake mushrooms and conventional electronics. Dehydration-based preservation was successfully explored, demonstrating the robustness of our devices. When used as RAM, our mushroom memristor was able to operate at up to 6000 Hz. Shiitake mushrooms are biodegradable and have demonstrated radiation resistance, suggesting that the potential applications of fungal computing range from sustainable computing devices to aerospace technologies.

## Acknowledgments

We would like to thank Ryan Lingo and Rajeev Chhajer of the Honda Research Institute.

## Author Contributions

J. L. provided the concept, project oversight, and manuscript writing. Q. T. provided resources and support. R. P. provided lab space to conduct the experiment. The direct investigation was conducted by J. S. and J. H.

## Data availability

Data is available upon request from the corresponding author.

## Additional Information

Supplementary Information is available for this paper.

Correspondence and requests for materials should be addressed to John LaRocco.

